# Creation and Validation of the First Infinium DNA Methylation Array for the Human Imprintome

**DOI:** 10.1101/2024.01.15.575646

**Authors:** Natalia Carreras-Gallo, Varun B. Dwaraka, Dereje D. Jima, David A. Skaar, Tavis L. Mendez, Antonio Planchart, Wanding Zhou, Randy L. Jirtle, Ryan Smith, Cathrine Hoyo

**Affiliations:** TruDiagnostic, Inc., Lexington, KY USA; Center for Human Health and the Environment, North Carolina State University, Raleigh, NC, USA; Bioinformatics Research Center, North Carolina State University, Raleigh, NC, USA; Toxicology Program, North Carolina State University, Raleigh, NC, USA; Department of Biological Sciences, North Carolina State University, Raleigh, NC, USA; Center for Computational and Genomic Medicine, Children’s Hospital of Philadelphia, Philadelphia, PA, USA; Department of Pathology and Laboratory Medicine, University of Pennsylvania, Philadelphia, PA, USA

**Keywords:** Genomic imprinting, Imprintome, Imprint Control Region (ICR), Custom methylation array, DNA methylation

## Abstract

**Background:** Differentially methylated imprint control regions (ICRs) regulate the monoallelic expression of imprinted genes. Their epigenetic dysregulation by environmental exposures throughout life results in the formation of common chronic diseases. Unfortunately, existing Infinium methylation arrays lack the ability to profile these regions adequately. Whole genome bisulfite sequencing (WGBS) is the unique method able to profile these regions, but it is very expensive and it requires not only a high coverage but it is also computationally intensive to assess those regions.

**Findings:** To address this deficiency, we developed a custom methylation array containing 22,819 probes. Among them, 9,757 probes map to 1,088 out of the 1,488 candidate ICRs recently described. To assess the performance of the array, we created matched samples processed with the Human Imprintome array and WGBS, which is the current standard method for assessing the methylation of the Human Imprintome. We compared the methylation levels from the shared CpG sites and obtained a mean R^2^ = 0.569. We also created matched samples processed with the Human Imprintome array and the Infinium Methylation EPIC v2 array and obtained a mean R^2^ = 0.796. Furthermore, replication experiments demonstrated high reliability (ICC: 0.799-0.945).

**Conclusions:** Our custom array will be useful for replicable and accurate assessment, mechanistic insight, and targeted investigation of ICRs. This tool should accelerate the discovery of ICRs associated with a wide range of diseases and exposures, and advance our understanding of genomic imprinting and its relevance in development and disease formation throughout the life course.

## Introduction

Interdisciplinary evidence accumulated over the past decades supports that environmental stressors — where the environment is broadly defined to include the built environment, contaminants, and psychosocial and socioeconomic stressors —are estimated to contribute 70– 90% of the burden of common chronic diseases in humans [1–3]. Nevertheless, empirical human data remains limited as disease formation often occurs many years after environmental exposure.

Epigenetics, a means by which genes respond to a wide range of environmental stressors to stably change gene expression, replication, and repair can cause long-term health effects [4]. Mechanistically, this is achieved by altering the organization and function of chromatin with the use of modifications to DNA and histones, as well as non-coding RNA interference [5]. The most studied epigenetic mechanism in humans is DNA methylation, owing to the stability of the DNA molecule. Indeed, data generated in the last decade, much of it from multiple iterations of Illumina Methylation Arrays, demonstrate the mediating role of CpG methylation in the effects of a wide range of environmental exposures and chronic diseases [6–8]. Sometimes the interpretation of the findings is ambiguous because the DNA methylation modifications identified in the accessible tissues, such as blood, are different from the marks in the diseased tissue [9, 10]. Moreover, many epigenetic marks must respond to environmental exposure throughout the life course, making cause-and-effect difficult to infer in related disease outcomes.

Known exceptions to this variability are CpG methylation marks that are stochastically established before tissue specification that controls metastable epiallele expression [11], and those that reside in the imprint control regions (ICRs) that regulate the monoallelic expression of imprinted genes [12–14]. CpG methylation in ICRs is established before gastrulation and is mitotically heritable, such that these epigenetic marks are normally similar across tissues and cell types over the life course. ICRs are defined by parent-of-origin specific methylation marks that are important gene dosage regulators and are similar across individuals. The stability of these methylation marks with age also makes them long-term ‘records’ of early exposures that are difficult to obtain through questionnaires or other exposure assessment assays [14].

Moreover, multiple lines of evidence support that changes in the methylation patterns in ICRs are implicated in many disorders such as cancer [15, 16], neurological disorders [17], specific syndromes such as Prader–Willi syndrome [18] and Angelman syndrome [19], as well as chronic diseases that result from exposure to environmental contaminants [20, 21].

While these features make ICRs attractive targets for unraveling the mechanisms underlying many chronic diseases with developmental origins and in developing potential therapeutic strategies, until a year ago, only 24 ICRs were characterized [13, 22]. In a recent study, Jima *et al*. identified 1,488 candidate differentially methylated ICRs in humans by performing whole-genome bisulfite sequencing (WGBS) [12]. The replicable measurement of CpG methylation in these regions in a large number of samples is only possible using a high throughput DNA methylation sequencing method.

The most common DNA methylation microarrays in academic and commercial research are the Illumina Infinium HumanMethylation450, the Infinium Illumina EPIC850k BeadChip, and the Infinium Methylation EPICv2 BeadChip (Illumina, Inc., San Diego, CA). Among the 22,408 CpG sites mapped to the 1,488 ICRs, only 6.7% of these probes are represented on the EPICv2 array. Thus, the current arrays are not able to comprehensively profile ICRs, limiting our ability to use these arrays to investigate the role of imprinting in disease development.

Here we describe the development of a custom methylation array specifically designed to target and measure the methylation status of the candidate ICRs involved in imprinted gene expression. This technology provides a targeted and comprehensive approach that allows accurate assessment, mechanistic insights, and the identification of aberrations within ICRs associated with various diseases.

## Methods

### Sample Cohort information

#### Alzheimer’s disease brain tissues

DNA derived from autopsy brain specimens was obtained from the Joseph and Kathleen Bryan Brain Bank of the Duke University/University of North Carolina at Chapel Hill Alzheimer’s Disease Research Center (Duke/UNC ADRC). These brain tissues were selected according to their neuropathologic diagnosis of Alzheimer’s disease. Eight brain samples from AD autopsies (4 non-Hispanic Blacks – NHBs - and 4 non-Hispanic Whites – NHWs) and eight brain samples from control autopsies (4 NHBs and 4 NHWs) were processed using both the Human Imprintome array and WGBS with 10-15X coverage (paper in review).

#### Alzheimer’s disease whole blood

A pilot case control study of 50 cases and 50 controls was also conducted at the Duke Memory Clinic. Peripheral whole blood was collected by the lancet and capillary method into lysis buffer and DNA extracted. In total, DNA samples from 17 individuals were randomly selected. Among them, 10 were Alzheimer’s disease cases and 7 were controls. All the samples were processed twice using the Human Imprintome array to assess the performance of the array. Moreover, three more controls were randomly selected to process them with the Human Imprintome array and the EPICv2 array.

#### Newborn epigenetics study (NEST) cohort umbilical cord

NEST is an ongoing prospective birth cohort study with 2,681 pregnant women recruited in two waves between 2005 and 2011; enrollment of participants is described in detail elsewhere [23, 24]. Briefly, pregnant women were recruited from prenatal clinics serving Duke University Hospital and Durham Regional Hospital obstetrics facilities in Durham, North Carolina. Eligible participants were: (1) pregnant, (2) at least 18 years of age, (3) English-speaking, and (4) intending to deliver at one of two obstetric facilities. Women with HIV or intending to give up custody of their offspring were excluded. At delivery, umbilical cord blood was obtained. For the current study, we used 8 umbilical cord blood samples, and processed them twice with the Human Imprintome Array to assess the reliability of the array.

### Preprocessing of the Human Imprintome and EPIC array data

For the preparation of samples, 200 ng of DNA were bisulfite converted using the EZ DNA Methylation kit (Zymo Research, Irvine, CA) according to the manufacturer’s instructions. Bisulfite-converted DNA samples were randomly assigned to a chip well on the Infinium Human Methylation EPIC v2 BeadChip (Illumina, Inc., San Diego, CA) or in the Human Imprintome array BeadChip (Illumina, Inc., San Diego, CA), amplified, hybridized onto the array, stained, washed, and imaged with the Illumina iScan SQ instrument (Illumina, Inc., San Diego, CA) to obtain raw image intensities. For the EPIC processing, the data were processed at TruDiagnostic (Inc., Lexington, KY) using a custom EPIC v2 array that include all the probes from the regular EPICv2 array [25] and 6,930 additional probes spiked in.

To preprocess the DNA methylation values, we utilized the *sesame* package [26] due to its compatibility with custom arrays. Specifically, we employed the *readIDATpair* followed by the *getBetas* functions, which require the IDAT file locations and the custom manifest to generate a beta value matrix. To visualize the distribution of beta values we employed the *densityPlot* function from the *minfi* package [27].

### Processing of the WGBS data

For the Alzheimer’s disease autopsy tissues (n=16), libraries were prepared using EpiGnome™ Methyl-Seq reagents (Illumina, Inc, *San Diego, CA*), index-tagged for multiplexing, and sequenced on an Illumina NextSeq platform (Illumina, Inc, *San Diego, CA*). Reads were assigned back to individuals by indexing, and aligned *in silico* to a bisulfite-converted reference genome (version Hg38), eliminating reads without unique alignments (due to either repetitive genomic sequence or loss of specificity from bisulfite conversion of cytosines) and duplicate reads (indicative of clonal amplification of original random DNA fragments). From these reads, methylation fractions and read counts were calculated for all CpG sites in the genome. There was >97% bisulfite conversion in all samples, with sequence coverage between 10X-15X and no sequence duplication bias. To compare against array data, the percent of methylated reads over the total number of reads was used to estimate overall methylation values. We removed the CpG sites that had fewer than 10 reads per probe from the WGBS data to ensure a correct estimation of the methylation level. These values were then compared to unnormalized beta values extracted from arrays.

## Results

### Design of the Human Imprintome array

For the development of the custom Human Imprintome array, we used the 22,408 probes that were mapped to the 1,488 ICRs described by Jima et al using WGBS [12] (Table S1). Previous studies have identified the necessity to exclude low-specificity probes that can bind to multiple sequences within the genome, as well as probes that contain genetic variants in their underlying sequence [28, 29]. Among the 22,408 CpG sites, we identified 1,313 multimapper probes. The number of mapping genomic positions per probe in the GRCh38 Build Genome ranged from 2 to 100 (Fig. S1; Table S2). Since these 1,313 multimapper probes were representative of a high number of ICRs, we decided to keep these probes on the array although the chromosome and the position in the manifest were set to 0 to avoid confusion.

Second, we used threshold scores designed and validated by Illumina, Inc. (San Diego, CA) to only select high-quality probes. Mainly, we used a general score based on probe sequence, including GC content and annealing temperature. The minimum probe score was set to 0.3 for the converted strand and 0.2 for the opposite strand. Lastly, we included control probes in the array for normalization purposes.

As a result, the Human Imprintome array manifest (Table S3) comprises a total of 22,819 probes, categorized into 704 control probes and 22,115 CG probes (Table 1). Out of the CG probes, 9,757 were successfully aligned with one of the 1,488 identified ICRs (Table S4). The remaining CG probes served various purposes, including multimapping, background normalization, or mapping to distinct locations that did not intersect with any ICR. The manifest includes 10,364”cgBK” probes with missing chromosomes that were included in the manifest to enable any additional background-normalization capabilities. Among them, there are 9,163 Infinium II assays, 485 Green extending Infinium I probes, and 716 Red extending Infinium I probes.

**Table 1.** Description of the probes contained in the Human Imprintome Array.

| Probe Type | # Probes | Description |
| --- | --- | --- |
| CG - Unique mappers | 10,438 | Probes mapping to unique locations. 9,757 probes of them are mapping to ICRs. |
| CG - Background normalization | 10,364 | Probes with "cgBK" prefix included in the manifest to enable additional background normalization capabilities |
| CG - Multimapping | 1,313 | Probes mapping to multiple locations |
| Bisulfite conversion I | 10 | Infinium I probes to monitor efficiency of bisulfite conversion |
| Bisulfite conversion II | 4 | Infinium II probes to monitor efficiency of bisulfite conversion |
| Extension | 4 | Control probes to test the efficacy of A, T, C, and G nucleotides from a hairpin probe |
| Hybridization | 3 | Control probes to test the overall performance of the entire assay using synthetic targets instead of amplified DNA |
| Negative | 411 | Control probes that should not hybridize to the DNA template |
| Non-polymorphic | 9 | Non-polymorphic control probes |
| Norm_A | 27 | Internal normalization control probes that incorporate a base A |
| Norm_C | 58 | Internal normalization control probes that incorporate a base C |
| Norm_G | 27 | Internal normalization control probes that incorporate a base G |
| Norm_T | 58 | Internal normalization control probes that incorporate a base T |
| Restoration | 1 | Control probes to assess the efficiency of DNA restoration in the Infinium HD FFPE protocol<sup>a</sup> |
| rs | 69 | Single Nucleotide Polymorphism control probes |
| Specificity I | 12 | Control probes to monitor allele-specific extension for Infinium I probes |
| Specificity II | 3 | Control probes to monitor allele-specific extension for Infinium II probes |
| Staining | 6 | Control probes to examine the efficiency of the staining step in both the red and green channels |
| Target removal | 2 | Control probes to test the efficiency of the stripping step after the extension reaction |

<sup>a</sup> Restoration probes are only informative when following the Infinium Formalin-Fixed Paraffin-Embedded (FFPE) workflow.

Illumina, Inc. (San Diego, CA) could not guarantee the inclusion of all the probes in the final array design. Thus, out of the 1,488 ICRs, a subset of 1,088 ICRs (73.1%) had successful probe alignments, as outlined in Table 2. The distribution of probes per ICR in the Human Imprintome array revealed a mean value of 9, encompassing a range from a minimum of 1 probe to a maximum of 171 probes. Notably, a significant proportion (n = 672) of ICRs exhibited successful mappings with more than 5 probes.

**Table 2.**
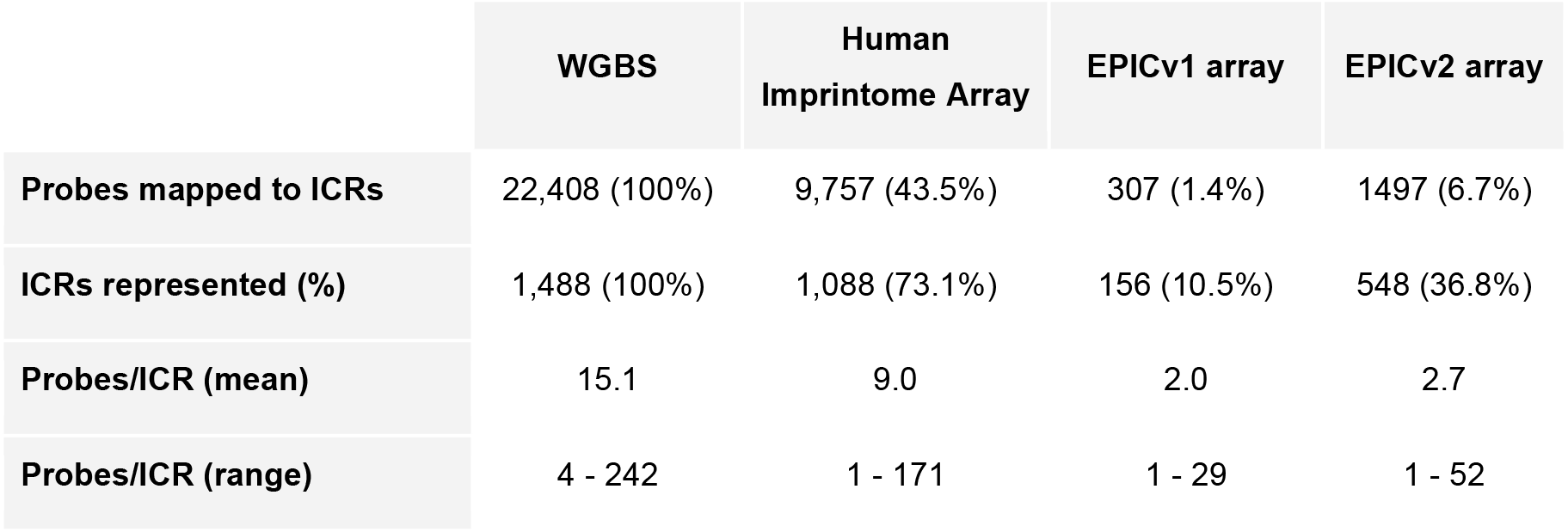
Representation of the Human Imprintome in the Human Imprintome array, EPICv1 array, and EPICv2 array compared to the current standard method, Whole genome bisulfite sequencing (WGBS).

|  | WGBS | Human Imprintome Array | EPICv1 array | EPICv2 array |
| --- | --- | --- | --- | --- |
| Probes mapped to ICRs | 22,408 (100%) | 9,757 (43.5%) | 307 (1.4%) | 1497 (6.7%) |
| ICRs represented (%) | 1,488 (100%) | 1,088 (73.1%) | 156 (10.5%) | 548 (36.8%) |
| Probes/ICR (mean) | 15.1 | 9.0 | 2.0 | 2.7 |
| Probes/ICR (range) | 4 - 242 | 1 - 171 | 1 - 29 | 1 - 52 |

### Comparison with other arrays

To compare the Human Imprintome array with other arrays, we calculated the ICR representation in each sequencing method. As a reference, we used the WGBS method and the 22,408 CpG sites that were identified to be representative of the 1,488 ICRs [12]. Fig. 1 demonstrates that the Imprintome array has a much larger representation of ICRs compared to EPICv1 and EPICv2 arrays. As shown in Table 2, the EPICv1 array only examines 307 probes out of the 22,408 (1.4%), and has a representation of 156 ICRs. Moreover, the average number of probes per ICR is 2. The last version of the EPIC array, version 2, can sequence a higher number of Human Imprintome probes compared to version 1. In this case, 6.7% of the probes are analyzed and they represent 548 ICRs with an average of 2.7 probes per ICR. In contrast, the Human Imprintome array evaluates the methylation level of 9,757 probes that are mapped to unique locations (43.5%), and represent 1,088 ICRs with an average of 9 probes per ICR.

**Fig. 1.**
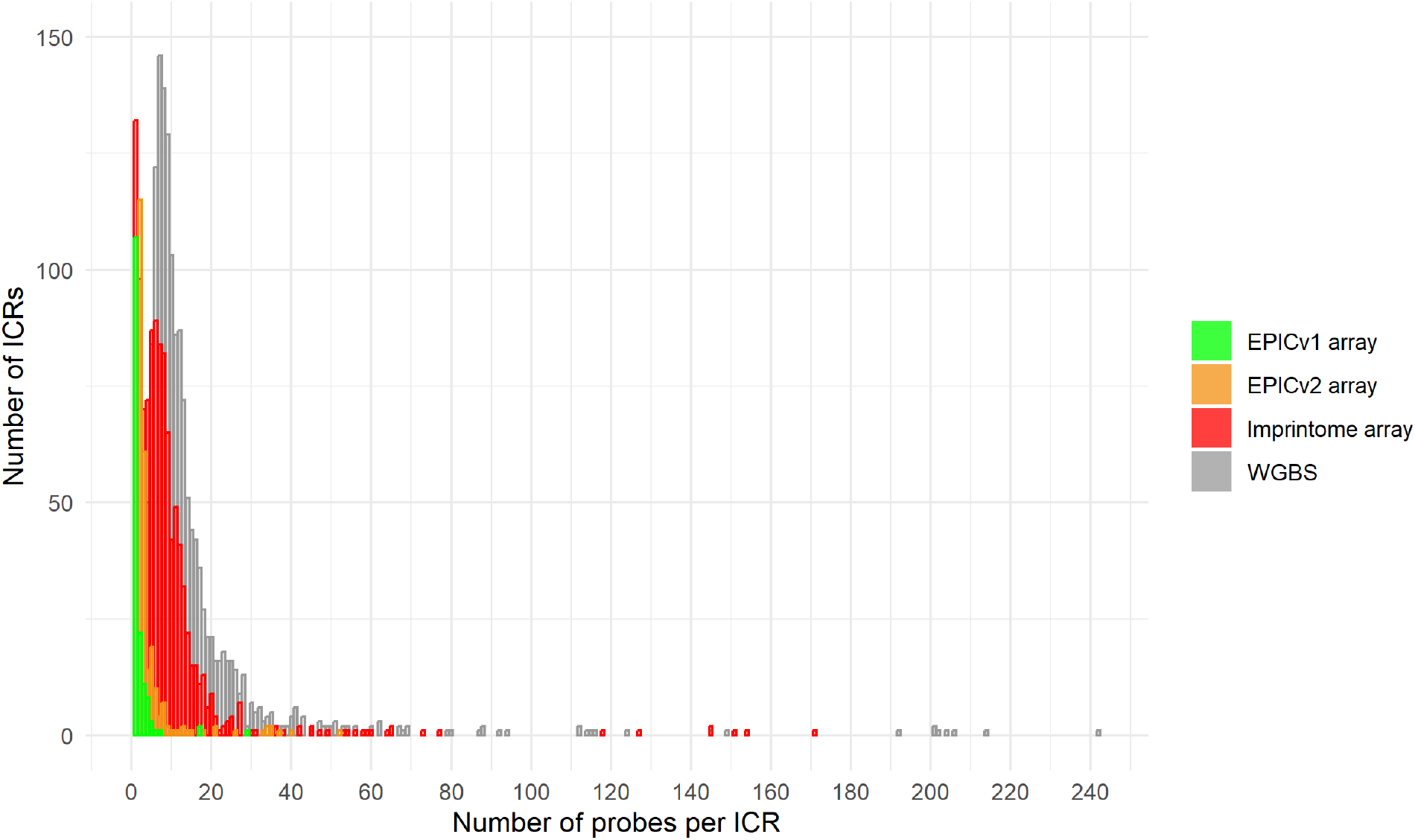
Histogram for the number of probes mapped to each ICR for each sequencing method. Whole genome bisulfite sequencing (WGBS) is the reference and it contains the 1488 ICRs described, as well as the 22,408 probes mapped to these regions. The other sequencing methods have a lower number of ICRs and probes representing those ICRs.

To check whether the beta values obtained using the Human Imprintome array are comparable with the beta values using the EPICv2 array, we examined three whole blood samples from the Alzheimer’s cohort that were processed using both the EPICv2 array and the Human Imprintome array. It is important to note that while the methylation levels from the EPIC array and WGBS method are derived from CpG sites, the beta values from the Human

Imprintome array correspond to probes. Consequently, a single CpG site may be represented by multiple probes on the Human Imprintome array. To ensure comparability across arrays, we collapsed multiple probes targeting the same CpG site by calculating their mean.

Utilizing this approach, we identified 1,703 probes sites that overlapped between the EPICv2 and Human Imprintome arrays. We obtained Pearson correlation coefficients of 0.788, 0.811, and 0.789, respectively, as shown in Fig. 2.

**Fig. 2.**
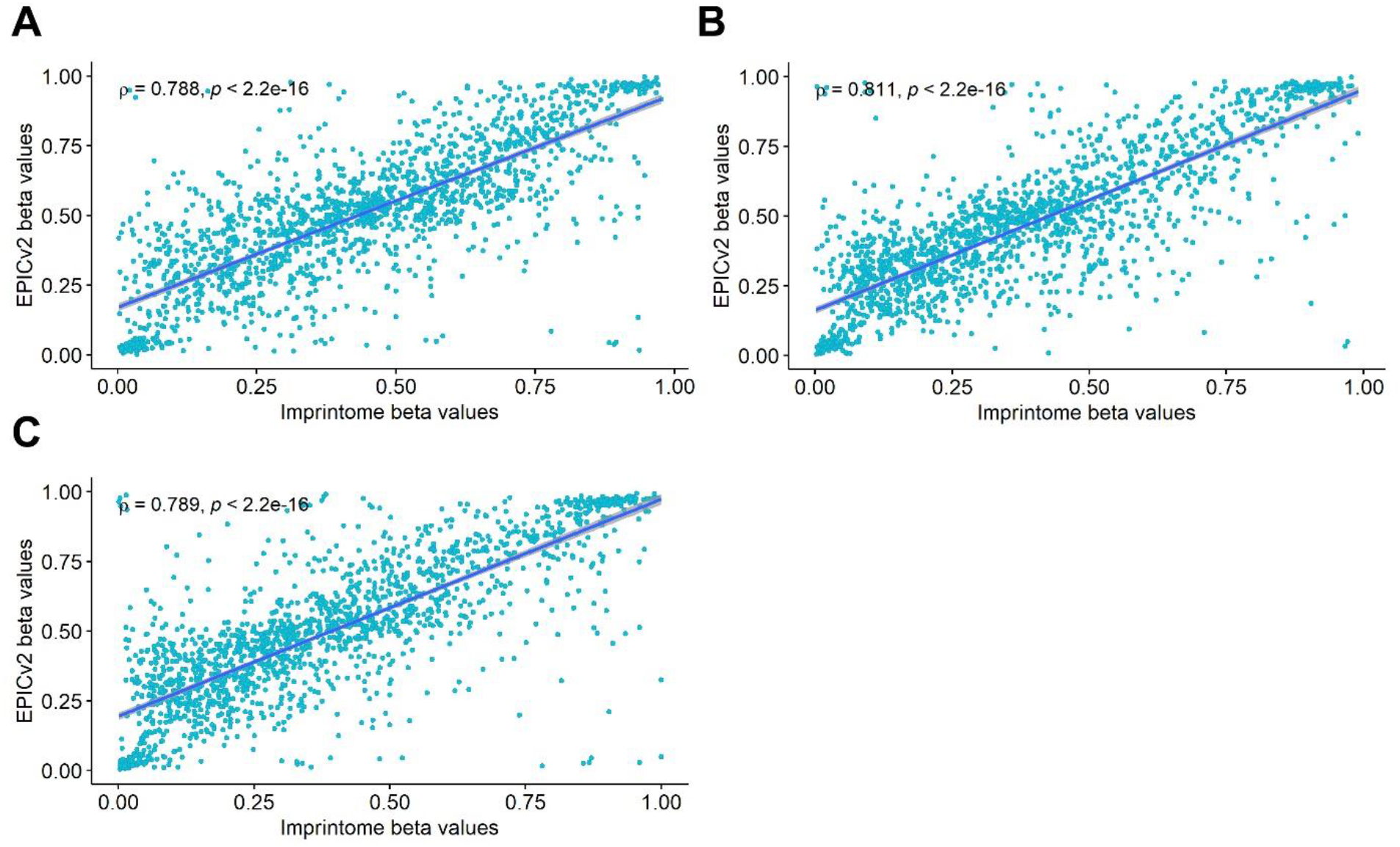
Correlation plots between Human Imprintome and the Infinium Methylation EPIC arrays. We selected the 1703 probes overlapped between both arrays to plot the Pearson correlation. (**A**) Sample AD-125, which belongs to a female Non-Hispanic White (NHW) sample. (**B**) Sample AD-157, which belongs to a male Non-Hispanic White sample. (**C**) Sample AD-173, which belongs to a male Non-Hispanic Black (NHB) sample.

Moreover, using the same samples, we compared the beta value density plots using all the probes from each array. The beta values in the Human Imprintome array exhibit a peak frequency at 0.5, which is indicative of monoallelic methylation (Fig. 3A). This pattern arises from the presence of approximately 100% methylation on one parental allele and 0% methylation on the other. The density plot of the probes analyzed in the EPICv2 array displays two peaks at 0 and 1, reflecting biallelic methylation, where both probes were either methylated or unmethylated (Fig. 3B). This shows that the Human Imprintome array is correctly assessing the methylation levels of the Human Imprintome probes.

**Fig. 3.**
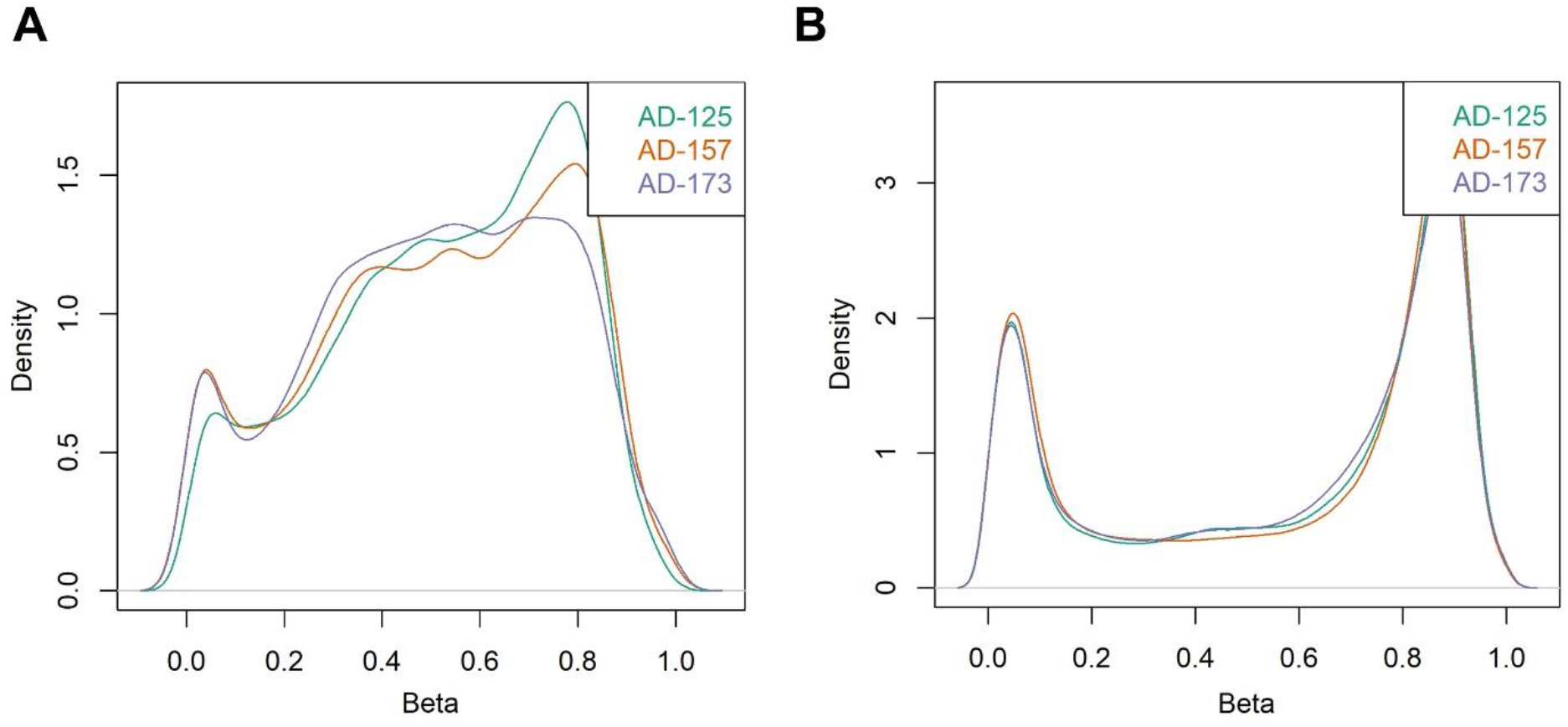
Comparison of the beta distribution between the Human Imprintome array (**A**) and the Infinium Methylation EPIC v2 array (**B**) in the same samples. The beta values in the Human Imprintome array exhibit a peak frequency at 0.5, which is indicative of monoallelic methylation. This pattern arises from the presence of approximately 100% methylation on one parental allele and 0% methylation on the other. The density plot of the probes analyzed in the EPICv2 array displays two peaks at 0 and 1, reflecting biallelic methylation, where both probes were either methylated or unmethylated.

Additionally, we compared the Human Imprintome array to the current standard method for assessing the methylation of the Human Imprintome (i.e., WGBS). We used 16 brain samples from Alzheimer’s cases and controls. These samples were classified into 4 controls and 4 cases of NHBs and 4 controls and 4 cases of NHWs. To correctly assess the correlations between the Human Imprintome array and WGBS, we combined the 4 samples from each group. This increased the coverage and obtained a higher number of shared CpG sites between both methodologies. We collapsed the probes that were targeting the same CpG site in the Human Imprintome array.

We removed the CpG sites that had fewer than 10 reads per probe from the WGBS data because the probability of estimating an incorrect methylation level is higher when the number of reads per probe is low. We then calculated the percentage of methylated reads compared to the total of reads mapped to each probe to compare this value to the beta values from the Human Imprintome array. Finally, we calculated the correlations using the CpG sites shared in each group. The highest number of CpG sites shared was for NHB cases with 7746 and the lowest was 4513 for NHW controls. The correlations ranged from 0.532 for NHW cases to 0.657 for NHB controls with a mean of 0.569 (Fig. 4). We also did a filtration at 1, 3, 5, and 20 reads per probe but the best results with a sufficient number of probes to compare were obtained with 10 (Table S5).

**Fig. 4.**
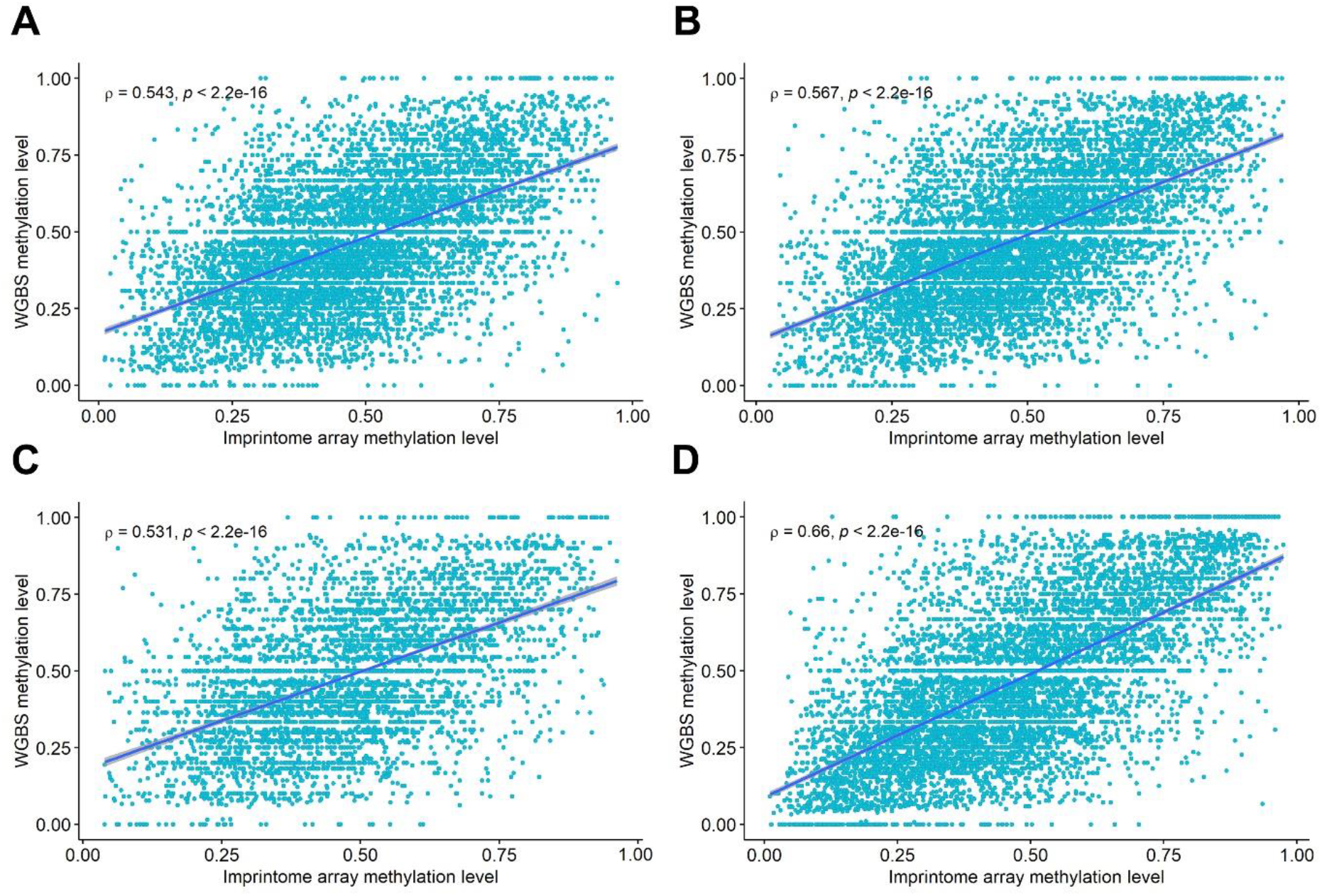
Correlation between the Human Imprintome array and WGBS. (**A**) Correlation plot for the combination of four Non-Hispanic White (NHW) Alzheimer’s Disease cases. (**B**) Correlation plot for the combination of four Non-Hispanic Black (NHB) Alzheimer’s Disease cases. (**C**) Correlation plot for the combination of four NHW controls. (**D**) Correlation plot for the combination of four NHB controls.

### Performance in replicates

To test the performance of the array, we analyzed replicates in the laboratory. We used 8 samples from the umbilical cord blood (NEST cohort [23,24]) and 17 samples from whole blood (Alzheimer’s cohort). Using all the probes, we calculated the intraclass correlation (ICC) between the first and the second replicate for each sample. The ICC values ranged from 0.799 to 0.945, having a mean of 0.868 (Table S6; Fig. S2-S3).

## Discussion

The Illumina BeadArray technology has undergone substantial redevelopment over the years, and the total number of CpG sites that can be simultaneously analyzed has increased substantially from ∼25,000 in 2008 (HumanMethylation27KBeadChip) [30], to ∼485,000 in 2011 (HumanMethylation450K BeadChip) [31], to over ∼850,000 CpG sites in 2016 (MethylationEPIC BeadChip v1.0) [32], and finally to over ∼935,000 CpG sites in 2022 when MethylationEPIC BeadChip v2.0 was released in June of 2023 [25].

In this study, we have successfully developed and characterized a Human Imprintome custom array, which offers an innovative approach for investigating DNA methylation patterns specifically associated with ICRs and parental allele-specific methylation. Using replicate samples, we have demonstrated that this custom array exhibits exceptional performance in accurately capturing DNA methylation information. Furthermore, the analysis of beta values obtained from the shared CpG sites between the Human Imprintome array, EPIC v2 array, and WGBS has revealed a high degree of correlation, reinforcing the robustness and reliability of the Human Imprintome custom array.

The Human Imprintome array represents a powerful tool that provides researchers with a focused platform for high-resolution analysis of DNA methylation dynamics in imprint control regions. Custom arrays like the Human Imprintome array can be a cost-effective solution compared to using commercially available arrays that contain probes unrelated to the research focus. By focusing on specific regions of interest, researchers can also optimize their resources, and obtain more relevant data. Notably, existing commercial arrays, such as the EPICv1 array, contain a limited number of probes mapping to ICRs, representing only 10.5% of the total ICRs.

Infinium arrays are extensively used and there are many bioinformatic pipelines to process them. Although not all of them work with custom arrays, *sesame* is an existing package that seamlessly integrates with the Human Imprintome array, and ensures accurate data processing [26]. Moreover, the utilization of arrays provides precise estimates of DNA methylation at specific sites, enabling the design of epigenetic biomarkers specific to the Human Imprintome array.

To enhance the utility of the Human Imprintome array, we have provided a comprehensive annotation of the probes contained in the array, as well as detailed information regarding the ICRs and the probes mapped to them. This annotation allows researchers to gain a deeper understanding of the regions and probes interrogated by the array, facilitating more targeted and insightful analyses.

Although the Human Imprintome array exhibits remarkable performance, we acknowledge its limitations, including the absence of probes representing 400 ICRs and the relatively low coverage in some represented ICRs. However, we are actively addressing these limitations, and working towards the development of a new version of the array that includes probes mapped to the missing ICRs.

In conclusion, the Human Imprintome custom array has the potential to significantly contribute to the identification of CpG sites and ICRs with altered methylation levels. This, in turn, may provide valuable insights into the emergence and evolution of diseases associated with aberrant DNA methylation patterns involved in imprinted gene regulation. The Human Imprintome array, with its focused design, extensive annotation, and promising performance, holds great promise as a powerful tool for unraveling the complex relationship between DNA methylation, imprinting, and disease pathogenesis.

## Acknowledgments

We thank the Newborn Epigenetics Study (NEST) families for their participation. We also thank Dr. Andrew Lui of the Duke Memory Clinic for enrolling Alzheimer’s disease cases and controls.

## Author’s contributions

RS, CH, RLJ, and DAS conceived and designed the study. NCG, VBD, and DDJ performed the bioinformatic analyses. DAS and TLM analyzed the samples at the lab. SKK provided the cell culture samples. All authors drafted and edited the manuscript.

## Availability of data and materials

The data that support the findings of this study are available upon request to ensure the privacy of the participants. Please for data requests. Sequence data will be available at the dbGaP within 12 months.

## Declarations

### Ethics approval and consent to participate

The study protocol from the NEST and Alzheimer’s Disease study were approved by the Duke University Institutional Review Board. Autopsy specimens were exempted from IRB review by Duke and NC State University IRBs.

## References

1. Rappaport SM, Smith MT. Environment and Disease Risks. Science. 2010;330:460–1.

2. Lichtenstein P, Holm NV, Verkasalo PK, Iliadou A, Kaprio J, Koskenvuo M, et al. Environmental and heritable factors in the causation of cancer--analyses of cohorts of twins from Sweden, Denmark, and Finland. N Engl J Med. 2000;343:78–85.

3. Willett WC. Balancing life-style and genomics research for disease prevention. Science. 2002;296:695–8.

4. Ho S-M, Johnson A, Tarapore P, Janakiram V, Zhang X, Leung Y-K. Environmental Epigenetics and Its Implication on Disease Risk and Health Outcomes. ILAR J. 2012;53:289–305.

5. Handy DE, Castro R, Loscalzo J. Epigenetic Modifications: Basic Mechanisms and Role in Cardiovascular Disease. Circulation. 2011;123:2145–56.

6. Mannens MMAM, Lombardi MP, Alders M, Henneman P, Bliek J. Further Introduction of DNA Methylation (DNAm) Arrays in Regular Diagnostics. Front Genet. 2022;13:831452.

7. Baylin SB, Jones PA. A decade of exploring the cancer epigenome - biological and translational implications. Nat Rev Cancer. 2011;11:726–34.

8. Robertson KD. DNA methylation and human disease. Nat Rev Genet. 2005;6:597–610.

9. Zhang B, Zhou Y, Lin N, Lowdon RF, Hong C, Nagarajan RP, et al. Functional DNA methylation differences between tissues, cell types, and across individuals discovered using the M&M algorithm. Genome Res. 2013;23:1522–40.

10. Lokk K, Modhukur V, Rajashekar B, Märtens K, Mägi R, Kolde R, et al. DNA methylome profiling of human tissues identifies global and tissue-specific methylation patterns. Genome Biol. 2014;15:r54.

11. Kessler NJ, Waterland RA, Prentice AM, Silver MJ. Establishment of environmentally sensitive DNA methylation states in the very early human embryo. Sci Adv. 2018;4:eaat2624.

12. Jima DD, Skaar DA, Planchart A, Motsinger-Reif A, Cevik SE, Park SS, et al. Genomic map of candidate human imprint control regions: the imprintome. Epigenetics. 2022;17:1920–43.

13. Skaar DA, Li Y, Bernal AJ, Hoyo C, Murphy SK, Jirtle RL. The Human Imprintome: Regulatory Mechanisms, Methods of Ascertainment, and Roles in Disease Susceptibility. ILAR J. 2012;53:341–58.

14. Hoyo C, Murphy SK, Jirtle RL. Imprint regulatory elements as epigenetic biosensors of exposure in epidemiological studies. J Epidemiol Community Health. 2009;63:683–4.

15. Plass C, Soloway PD. DNA methylation, imprinting and cancer. Eur J Hum Genet. 2002;10:6–16.

16. Monk D. Deciphering the cancer imprintome. Brief Funct Genomics. 2010;9:329–39.

17. Ho-Shing O, Dulac C. Influences of genomic imprinting on brain function and behavior. Curr Opin Behav Sci. 2019;25:66–76.

18. Cassidy SB, Schwartz S, Miller JL, Driscoll DJ. Prader-Willi syndrome. Genet Med Off J Am Coll Med Genet. 2012;14:10–26.

19. Williams CA, Driscoll DJ, Dagli AI. Clinical and genetic aspects of Angelman syndrome. Genet Med Off J Am Coll Med Genet. 2010;12:385–95.

20. Pathak R, Feil R. Environmental Effects on Genomic Imprinting in Development and Disease. In: Patel VB, Preedy VR, editors. Handbook of Nutrition, Diet, and Epigenetics. Cham: Springer International Publishing; 2019. p. 3–23.

21. Cowley M, Skaar DA, Jima DD, Maguire RL, Hudson KM, Park SS, et al. Effects of Cadmium Exposure on DNA Methylation at Imprinting Control Regions and Genome-Wide in Mothers and Newborn Children. Environ Health Perspect. 2018;126:037003.

22. Luedi PP, Dietrich FS, Weidman JR, Bosko JM, Jirtle RL, Hartemink AJ. Computational and experimental identification of novel human imprinted genes. Genome Res. 2007;17:1723–30.

23. McCullough LE, Mendez MA, Miller EE, Murtha AP, Murphy SK, Hoyo C. Associations between prenatal physical activity, birth weight, and DNA methylation at genomically imprinted domains in a multiethnic newborn cohort. Epigenetics. 2015;10:597–606.

24. Hoyo C, Murtha AP, Schildkraut JM, Forman MR, Calingaert B, Demark-Wahnefried W, et al. Folic acid supplementation before and during pregnancy in the Newborn Epigenetics STudy (NEST). BMC Public Health. 2011;11:46.

25. Kaur D, Lee SM, Goldberg D, Spix NJ, Hinoue T, Li H-T, et al. Comprehensive evaluation of the Infinium human MethylationEPIC v2 BeadChip. Epigenetics Commun. 2023;3:6.

26. Ding W, Kaur D, Horvath S, Zhou W. Comparative epigenome analysis using Infinium DNA methylation BeadChips. Brief Bioinform. 2023;24:bbac617.

27. Minfi: a flexible and comprehensive Bioconductor package for the analysis of Infinium DNA methylation microarrays | Bioinformatics | Oxford Academic. https://academic.oup.com/bioinformatics/article/30/10/1363/267584. Accessed 6 Sep 2023.

28. Hop PJ, Zwamborn RAJ, Hannon EJ, Dekker AM, van Eijk KR, Walker EM, et al. Crossreactive probes on Illumina DNA methylation arrays: a large study on ALS shows that a cautionary approach is warranted in interpreting epigenome-wide association studies. NAR Genomics Bioinforma. 2020;2:qaa105.

29. Chen Y, Lemire M, Choufani S, Butcher DT, Grafodatskaya D, Zanke BW, et al. Discovery of cross-reactive probes and polymorphic CpGs in the Illumina Infinium HumanMethylation450 microarray. Epigenetics. 2013;8:203–9.

30. Bibikova M, Le J, Barnes B, Saedinia-Melnyk S, Zhou L, Shen R, et al. Genome-wide DNA methylation profiling using Infinium® assay. Epigenomics. 2009;1:177–200.

31. Bibikova M, Barnes B, Tsan C, Ho V, Klotzle B, Le JM, et al. High density DNA methylation array with single CpG site resolution. Genomics. 2011;98:288–95.

32. Pidsley R, Zotenko E, Peters TJ, Lawrence MG, Risbridger GP, Molloy P, et al. Critical evaluation of the Illumina MethylationEPIC BeadChip microarray for whole-genome DNA methylation profiling. Genome Biol. 2016;17:208.

