## Additional File 1 for "Creation and Validation of the First Infinium DNA Methylation Array for the Human Imprintome"

Additional File 2

Natalia Carreras-Gallo[1], Varun B. Dwaraka[1], Dereje D. Jima[2,3], David A. Skaar[2,4,5], Tavis L. Mendez[1], Antonio Planchart[2,4,5], Wanding Zhou[6,7], Randy L. Jirtle[2,4,5], Ryan Smith[1], Cathrine Hoyo[2,4,5]

[1] TruDiagnostic, Inc., Lexington, KY USA

[2] Center for Human Health and the Environment, North Carolina State University, Raleigh, NC, USA

[3] Bioinformatics Research Center, North Carolina State University, Raleigh, NC, USA

[4] Toxicology Program, North Carolina State University, Raleigh, NC, USA

[5] Department of Biological Sciences, North Carolina State University, Raleigh, NC, USA

[6] Center for Computational and Genomic Medicine, Children's Hospital of Philadelphia, Philadelphia, PA, USA

[7] Department of Pathology and Laboratory Medicine, University of Pennsylvania, Philadelphia, PA, USA

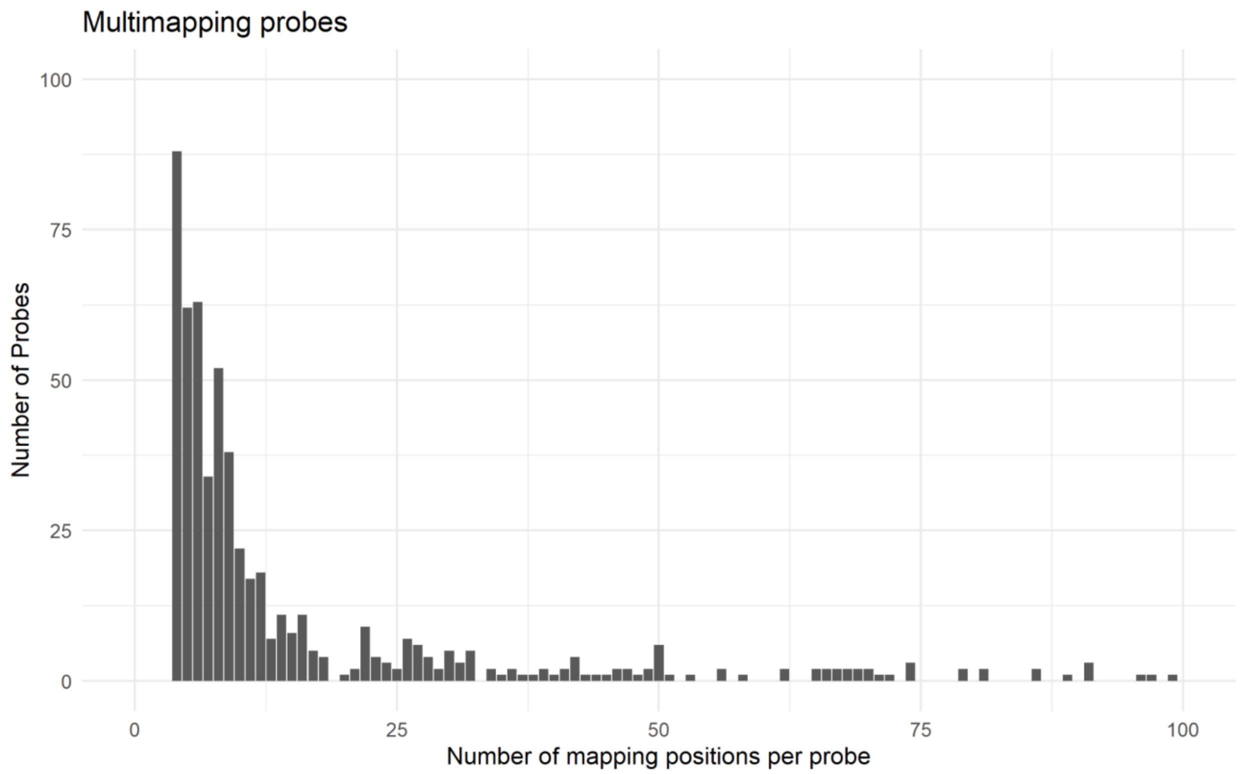


**Fig. S1** Number of mapping positions per multimapper probe in the Imprintome array. A total of 1,313 probes are multimapper and the minimum of genomic positions where they map is 2 and the maximum is 100.

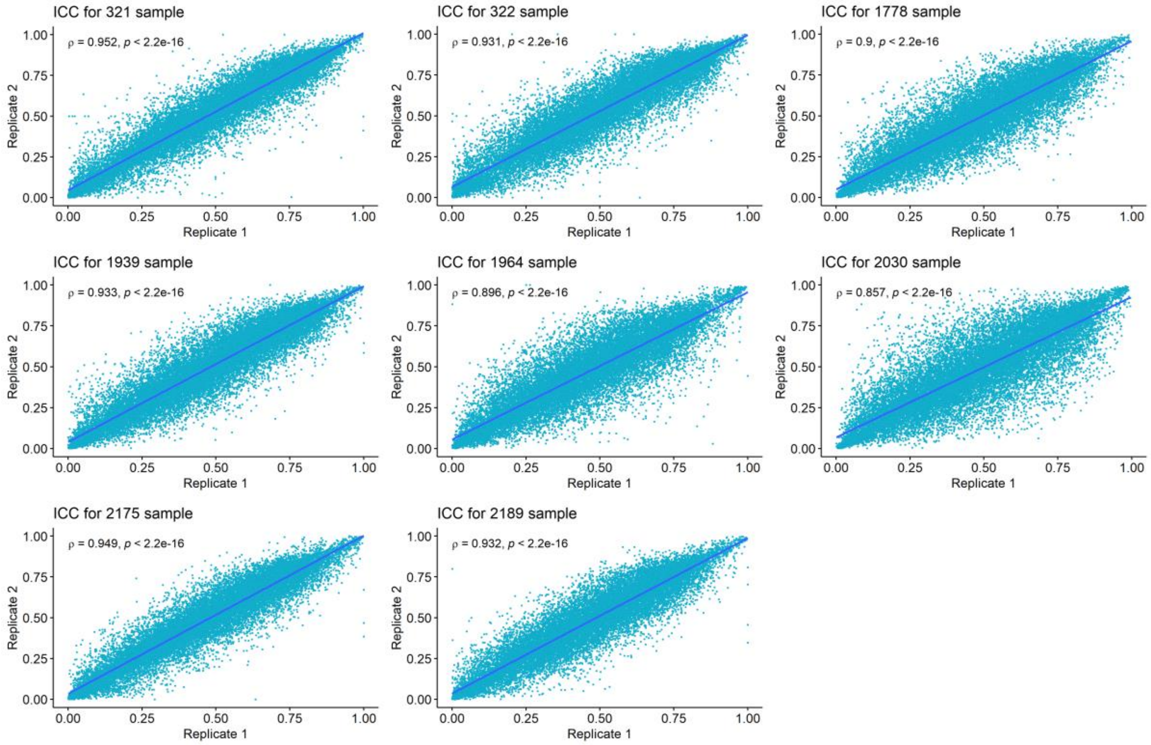


**Fig. S2** Intraclass correlation (ICC) values for 8 umbilical cord blood replicate samples.

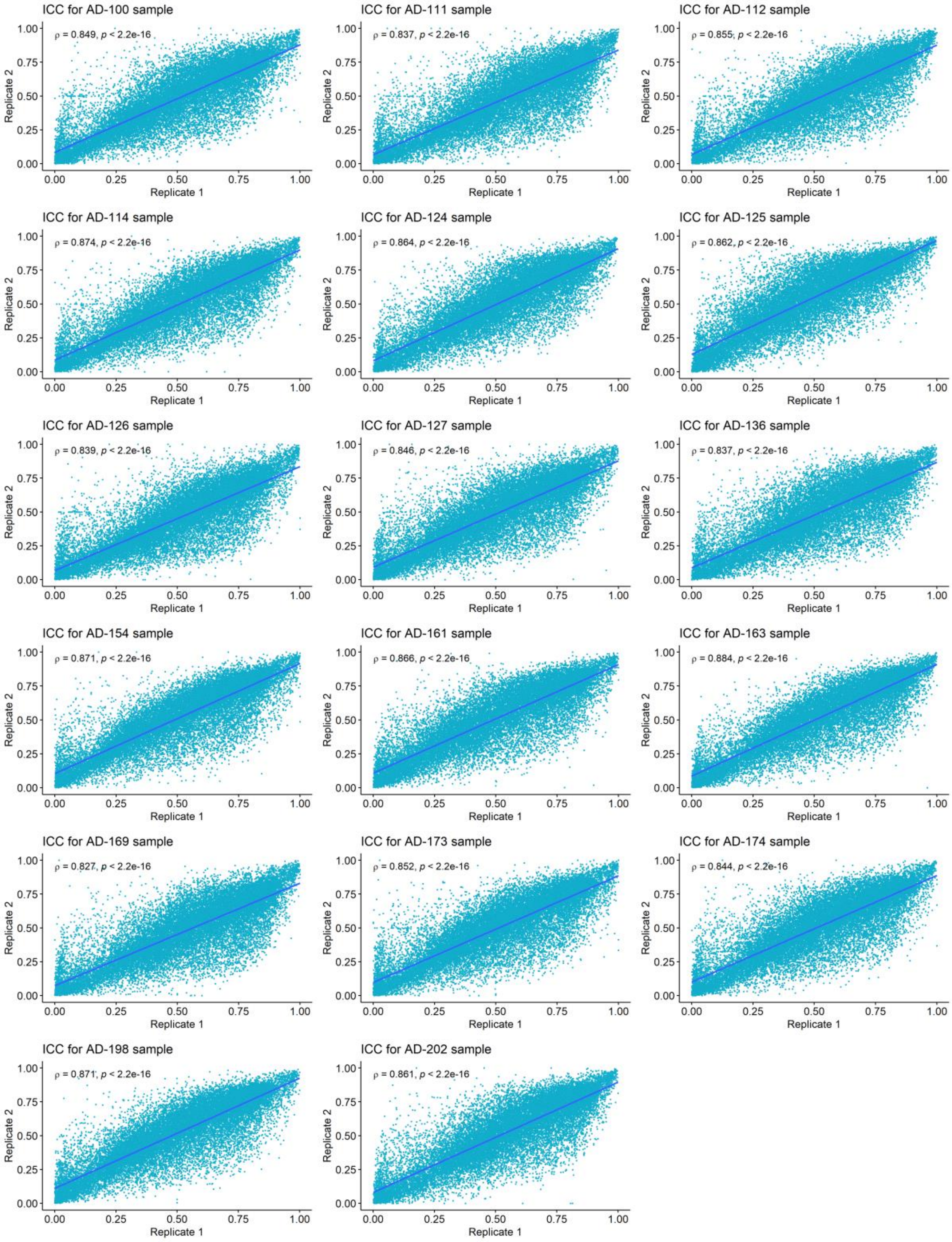


**Fig. S3** Intraclass correlation (ICC) values for 17 whole blood replicate samples.
